# Nonlinear functional responses and ecological pleiotropy alter the strength and likelihood of disruptive selection in consumers

**DOI:** 10.1101/283911

**Authors:** Kyle E. Coblentz

## Abstract

Much of the theory on disruptive selection has focused on selection in generalist consumers caused by ecological opportunity through the availability of alternative resources and intraspecific competition for those resources. This theory, however, makes several ecologically unrealistic assumptions. First, it assumes consumers have a linear, resource-dependent functional response, ignoring well-documented effects of resource handling times and consumer dependence. Second, it assumes the trait under selection only influences the per-capita attack rates of the consumer, ignoring other effects of the trait that may influence feeding rates and hence, fitness. Here, I develop a one consumer-two resource model to investigate how nonlinear functional responses and ecological pleiotropy (traits with multiple simultaneous ecological effects) influence the strength and likelihood of disruptive selection. I find that handling times and interference among consumers are capable of altering disruptive selection by changing feeding rates differentially across consumer phenotypes. In particular, handling times decrease the strength and likelihood of disruptive selection while the effects of interference depend on the mechanism through which interference occurs. The effects of handling times and interference, however, depend on whether and how ecological pleiotropy causes correlations between handling times or interference rates and attack rates. Overall, my results suggest that features underlying functional responses of consumers and the relationships among those features determine the likelihood and strength of disruptive selection. In particular, disruptive selection should be strongest in generalist populations with individuals who experience lower handling times and interference rates on the resources for which their attack rates are highest.

## Introduction

Disruptive selection, a process in which natural selection favors individuals with more extreme phenotypes over individuals with intermediate phenotypes, plays important roles in evolution and ecology. Evolutionarily, disruptive selection can cause and maintain genetic and phenotypic variation within populations and can drive speciation and adaptive diversification (Dieckmann and Doebeli, 1999; Doebeli, 2011; Rueffler et al., 2006; Smith, 1962). Ecologically, the intraspecific variation generated by disruptive selection can alter the interactions among species, their coexistence, and their functioning in ecosytems (Barbour et al., 2016; Bolnick et al., 2011; Gibert and Brassil, 2014; Gibert and DeLong, 2017; Hart et al., 2016; Hughes et al., 2008; Schreiber et al., 2011; Svanbäck et al., 2015).

Of the mechanisms generating disruptive selection, much of our knowledge comes from disruptive selection on resource-use traits in generalist consumers. Generally, disruptive selection on resource-use traits is thought to be a product of ecological opportunity through the availability of alternative resources, and intraspecific competition for those resources (Abrams et al., 2008; Nosil, 2012; Schluter, 2000). First, the availability of alternative resources provides a basis for fitness differences among individuals. If individuals with different traits are better able to use different resources, then individuals with phenotypes better matched to available resources will have greater fitness than individuals with intermediate phenotypes. If the mean trait of the consumer population lies between the optima for using the different available resources, disruptive selection will occur as individuals with more extreme phenotypes will have greater fitness than intermediate individuals. Intraspecific competition is then capable of stabilizing this form of disruptive selection by causing the selection to be frequency dependent. Given that different trait values affect individuals’ abilities to use different resources, the most common phenotypes will reduce the abundance of their associated resources to the greatest extent. The reduced availability of resources then drives higher rates of intraspecific competition in those common phenotypes. Consumers with less common phenotypes reduce their associated resources to a lesser extent and, all else being equal, experience less competition. The resultant increase in fitness for less common phenotypes leads to negative frequency-dependent disruptive selection because the relative fitness of phenotypes is dependent on their relative abundance in the population (Dieckmann and Doebeli, 1999; Smith, 1962).

Several studies have provided convincing empirical evidence that ecological opportunity through alternative resources and intraspecific competition cause disruptive selection in both the laboratory and field (Bolnick, 2001, 2004; Hendry et al., 2009; Martin and Pfennig, 2009). Yet, previous studies also suggest that the existence and strength of disruptive selection through these mechanisms are dependent on ecological features determining the relative availability of resources and the strength of intraspecific competition. For example, in a survey of fitness landscapes across populations of three-spined stickleback (*Gasterosteus aculeatus*), Bolnick and Lau (2008) showed that differences among populations in the existence and strength of disruptive selection was partially attributable differences in ecological opportunity among populations through differences in the relative availability of benthic versus limnetic habitat. In another survey of fitness landscapes, Martin and Pfennig (2012), showed that the strength of disruptive selection in spadefoot toad tadpoles (*Spea multiplicata*) was associated with the density of con-specifics, a proxy for the strength of intraspecific competition across populations. Together these results suggest that predicting the strength and occurrence of disruptive selection in consumers requires theory incorporating common ecological factors likely to influence ecological opportunity and intraspecific competition.

One factor likely to influence both ecological opportunity and the strength of intraspecific competition is the strength of the underlying consumer-resource interactions (Abrams et al., 2008; Jones and Post, 2013, 2016). For example, for intraspecific competition to influence disruptive selection consumers must deplete resources to an extent that it alters the fitness landscape across phenotypes, but, if the consumer-resource interactions are weak, prey depletion will be minimal and the resulting strength of disruptive selection will be weak (Abrams et al., 2008; Jones and Post, 2013,2016). Alternatively, if species interactions are strong, consumers can lead resources to local extinction, causing a decrease in ecological opportunity, again, altering strength of disruptive selection (Abrams et al., 2008; Jones and Post, 2013, 2016). One determinant of the strength of consumer-resource interactions largely ignored in current theory on disruptive selection in generalists is the consumer functional response. The consumer functional response defines the relationship between the densities of interacting species and consumer feeding rates and therefore is directly related to the strength of consumer-resource interactions and likely to influence disruptive selection. The vast majority of theory on disruptive selection in consumers assumes that the consumers have a linear, resource dependent functional response (Abrams et al., 2008; Ackermann and Doebeli, 2004; Dieckmann and Doebeli, 1999; Doebeli, 1978; Lawlor and Smith, 1976; MacArthur, 1972). However, linear functional responses are known to be rare (Jeschke et al., 2004). Nonlinearities in functional responses are the product of nearly ubiquitous properties of consumer-resource interactions such as handling times and consumer interference or facilitation (Abrams and Ginzburg, 2000; DeLong and Vasseur, 2011; Holling, 1959; Novak et al., 2017). Given the widespread nature of nonlinear functional responses and their effects on the strength of consumer-resource interactions, incorporating nonlinear functional responses into theory on disruptive selection in consumers may provide some insight into the characteristics of consumers most likely to exhibit disruptive selection.

Incorporating nonlinear functional responses into models of disruptive selection also provides an opportunity to address another assumption of most models: the traits of individuals only influence their per capita attack rates on resources. It is more likely that traits influencing an individual’s attack rates also influence other parameters of the functional response (i.e. handling and interference) causing parameters to be correlated across individuals. The presence of such correlated ecological trait effects has been termed ‘ecological pleiotropy’ (DeLong, 2017; DeLong and Gibert, 2016; Strauss and Irwin, 2004). Pleiotropic trait effects are likely to be common in functional responses. For example, a study of protists has shown attack rates, handling times, and interference rates to all covary with one another (DeLong, 2017). In another study, body size has been shown to have allometric relationships with both attack rates and handling times of consumers (Vucic-Pestic et al., 2009). Although ecologically pleiotropic trait effects in functional responses have been shown to alter population dynamics (DeLong, 2017) and have examined the evolutionary effects of pleiotropy between functional and numerical responses (Schreiber et al., 2018), it remains unclear how ecological pleiotropy may constrain or promote selection on underlying traits.

Here, I use a one consumer-two resource model to investigate how the parameters of nonlinear functional responses – attack rates, handling times, and interference rates – and potential correlations among them arising from an ecologically pleiotropic trait, influence the existence and strength of disruptive selection. My analyses indicate that nonlinear functional responses alter disruptive selection in ways that are dependent on the correlations between attack rates and handling times or attack rates and interference rates. This suggests that nonlinear functional responses and the presence of ecological pleiotropy can alter disruptive selection generated by intraspecific competition and ecological opportunity, offering a potential explanation for variation in the existence and strength of disruptive selection within and among systems.

## Methods

Below, I first introduce the general one consumer-two resource model used to investigate the relationships between nonlinear functional responses, ecological pleiotropy, and the strength of disruptive selection. I then explain the methods used to analyze the model. Lastly, I introduce the particular nonlinear functional responses investigated and explain how I modeled ecological pleiotropy among functional response parameters.

### The General Model

To investigate the effects of nonlinear functional responses and pleiotropy on the strength and likelihood of disruptive selection, I extended a one consumer-two resource model developed by Schreiber et al. (2011) to include nonlinear functional responses. The model begins with the assumption that the consumer population has a quantitative trait, *x*, that is normally distributed with mean, *x̄*, and variance, *σ*^2^, such that the distribution of *x* in the population, *p*(*x,x̄*), is described by, 
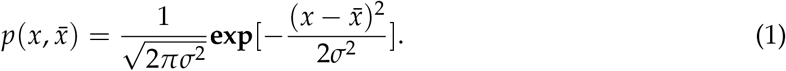

Furthermore, I assume that the phenotypic variance of the trait, *σ*^2^, consists of an environmental component, 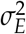, and a heritable genetic component, 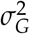, and that the heritable genetic component is positive. The value of an individual’s trait, *x*, is assumed to determine its attack rates, *α*_*i*_(*x*), on the two resources, *R*_*i*_ (*i* = 1,2), respectively. The maximum attack rate of an individual on the *i*th resource, *α*_*i,max*_, occurs at a trait value of *x* = *θ*_*i*_. The attack rate then decreases as the trait value moves away from *θ*_*i*_ in a Gaussian manner, 
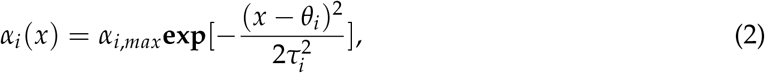
with the rate of decrease determined by 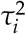. Letting *f*_*i*_ (*R*_1_, *R*_2_, *C*, *x*) denote the consumer’s functional response on resource *i* which depends on both resource densities, the trait *x*, and the consumer’s density in models including consumer interference, the mean fitness of the consumer population, 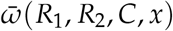, is: 
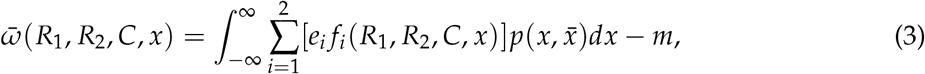
where *m* is the per capita mortality rate of the consumer and *e*_*i*_ is a linear conversion efficiency of resource *i* into consumers. Assuming logistic growth in the resources, the dynamics of the consumer and resource populations are: 
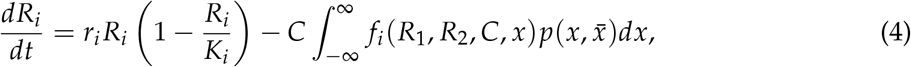

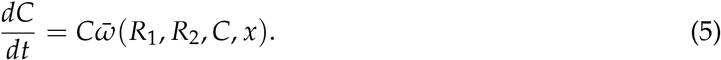
where *r*_*i*_ is the intrinsic growth rate of resource *i* and *K*_*i*_ is its carrying capacity.

Assuming that the consumer’s trait remains normally distributed and that the variance of the consumer’s trait remains constant and is not too large, the evolutionary dynamics of the mean of the consumer’s trait are described by: 
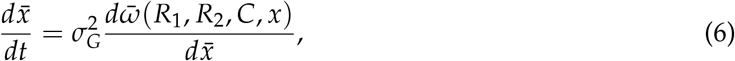

where 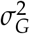 is the genetic component of the phenotypic variance of the consumer’s trait and 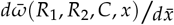 is the fitness gradient (Abrams et al., 1993; Iwasa et al., 1991; Lande, 1976). The fitness gradient describes directional selection on the mean of the consumer’s trait. At the ecological and evolutionary equilibrium of the system, there is no directional selection on the consumer’s trait and the fitness gradient is zero by definition. The resultant equilibrium is either a fitness maximum or a fitness minimum which can be determined from the curvature of the fitness function at the equilibrium given by the second derivative of the mean fitness with respect to consumer’s mean trait evaluated at the equilibrium, 
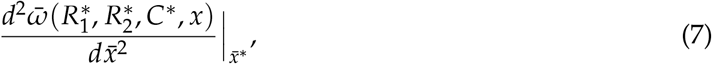
where the asterisks denote equilibrium values. If the curvature of the fitness function evaluated at the equilibrium is negative, the equilibrium is a fitness maximum and there is stabilizing selection on the trait. If the curvature of the fitness function is positive, the equilibrium is a fitness minimum and there is disruptive selection on the trait. The magnitude of the curvature of the fitness function provides a relative measure of the strength of selection. Thus, the curvature of the fitness function can be used to determine the parameters for which stabilizing or disruptive selection occur and the strength of that selection.

### Measuring Selection and Methods of Analysis

Here I consider a symmetric version of the above model in which all of the resource specific parameters are equal (e.g. *r*_1_ = *r*_2_, *K*_1_ = *K*_2_, etc.) except the *θ*_*i*_′s, and the *θ*_*i*_′s are symmetric about zero (i.e. *θ*_2_ = −*θ*_1_). Given the symmetric model, if the ecological dynamics reach a stable steady state, the evolutionary dynamics reach an equilibrium at *x̄*: = 0. The curvature of the fitness function at *x̄*: = 0 can then be used to determine the strength of selection and whether it is stabilizing or disruptive.

To determine how nonlinear functional responses and ecological pleiotropy alter the likelihood of disruptive selection, I compared numerical results of the model on the parameters at which selection changed from stabilizing to disruptive selection to analytical results derived by Schreiber et al. (2011). Using a symmetric version of the model with linear consumer functional responses, Schreiber et al. (2011) showed that disruptive selection occurs when, 
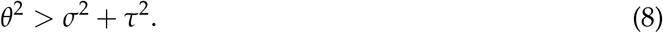

Therefore, if the consumers exhibited a linear functional response, selection switches from stabilizing to disruptive selection when *θ*^2^ = *σ*^2^ + *τ*^2^. I use this as a baseline to examine how nonlinear functional responses and ecological pleiotropy change the likelihood of disruptive selection relative to the case of consumers with linear functional responses. Specifically, I alter the value of *θ* while keeping *σ* and *τ* constant, although results are similar when varying *σ* and *τ*.

To evaluate how changes in the parameter values of nonlinear functional responses alter the strength and likelihood of disruptive selection, I performed numerical analyses of the model in Mathematica (v. 11.0.1.0, code for the analyses can be found in the Online Supplementary Material). I restricted my analyses to combinations of parameter values for which the consumer-resource interactions reached a fixed point equilibrium (i.e. did not exhibit cycles). After determining that the consumer-resource dynamics were at a fixed point using linear stability analysis, I varied the parameters of interest and calculated the curvature of the fitness function to determine whether selection was disruptive or stabilizing and the strength of selection. Although analytical results for the model were not possible, the results presented below were qualitatively similar across all of the different parameter values investigated that met the above criteria.

### Nonlinear Functional Response Due to Handling Times

To determine how handling times influence the strength and likelihood of disruptive selection, I substituted a multispecies Holling Type-II functional response (Holling, 1959), 
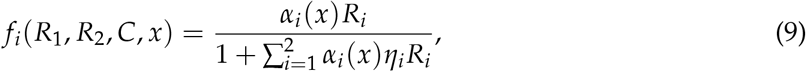
into equations 3-4, where *η*_*i*_ is the handling time of the consumer when feeding on resource *i*. I first investigated how handling times in general influenced the likelihood and strength of disruptive selection by assuming that the handling times for both resources were equal (i.e. *η*_1_ = *η*_2_). I then altered the magnitude of the handling times and examined the resulting effects on the likelihood strength of disruptive selection.

### Consumer-dependent Functional Responses

Although there are several ‘standard’ functional responses models that include interference, all were developed for a specialist consumer consuming a single resource (Beddington, 1975; Crowley and Martin, 1989; DeAngelis et al., 1975; Hassel and Varley, 1969). To examine the effects of interference on the likelihood and strength of disruptive selection in the one consumer-two resource model considered here, I extended two of the more mechanistic functional response models including interference – the Beddington-DeAngelis and Crowley-Martin functional responses – to more than one resource (Beddington, 1975; Crowley and Martin, 1989; DeAngelis et al., 1975).

The Beddington-DeAngelis and Crowley-Martin functional responses make different assumptions about how interference effects consumer feeding rates, and therefore could lead to differences in how they predict interference to alter the likelihood and strength of disruptive selection (Beddington, 1975; Crowley and Martin, 1989; DeAngelis et al., 1975). The Beddington-DeAngelis functional response model assumes that consumers interfere with one another at a rate, *γ*, and that interference decreases the time available for searching for resources. These assumptions lead to the following functional response for two resources, 
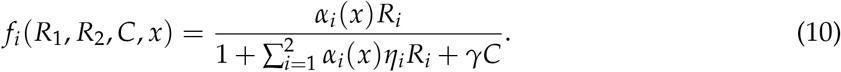

The Crowley-Martin functional response model extends the Beddington-DeAngelis functional response model by assuming that consumers also interfere while handling resources (Crowley and Martin, 1989). Under these assumptions, with a interference rate, *λ*, the Crowley-Martin functional response extended to two resources is, 
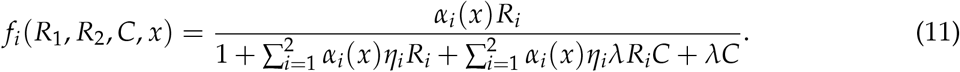

Given these two functional responses, I examined how interference in general changed the likelihood and strength of disruptive selection by altering the magnitude of interference and examining the resultant effects.

### Ecological Pleiotropy

To determine how an ecologically pleiotropic trait controlling both attack rates and handling times or attack rates and interference rates may influence the likelihood and strength of disruptive selection, I assumed that ecological pleiotropy causes attack rates and handling times or interference rates to covary.

For ecological pleiotropy affecting both attack rates and handling times, I modeled the effects of ecological plieotropy by making the handling times of each resource a linear function of the trait-dependent attack rates, 
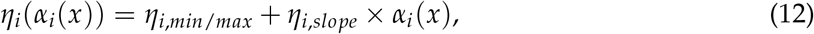
where *η*_*i*_(*α*_*i*_(*x*)) is the handling time of an individual on resource *i* with attack rate *α*_*i*_(*x*), *η*_*i,min/max*_ is the handling time of an individual on resource *i* when the individual has an attack rate of zero on resource *i*, and *η*_*i,slope*_ is the slope of the relationship exhibited across individuals with different trait values between the attack rates and handling times on resource *i*.

To model the effects of ecological pleiotropy affecting both attack rates and interference rates, I made the interference rate a linear function of the total attack rate of an individual on resources combined. Letting *y* represent either type of interference examined (i.e. *γ* for the Beddington-DeAngelis functional response or *λ* for the Crowley-Martin functional response), I modeled the correlation between interference and attack rates as, 
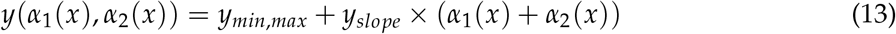
where *y*(*α*_1_ (*x*), *α*_2_ (*x*)) is the interference rate of an individual with attack rates *α*_1_(*x*) and *α*_2_ (*x*), *y*_*min,max*_ is the interference rate that would be experienced if both attack rates were zero, and *y*_*slope*_ is the slope of the relationship between the total attack rate interference rates across individuals having different trait values.

I considered both positive and negative correlations between attack rates and handling times and attack rates and interference rates. For positive correlations, *η*_*i,min/max*_ and *y*_*min/max*_ are minimums and *η*_*i,slope*_ and *y*_*slope*_ are positive. For negative correlations, *η*_*i,min/max*_ and *y*_*min/max*_ are maximums and *η*_*i,slope*_ and *y*_*slope*_ are negative. To examine the effects of the correlations between attack rates and handling times or attack rates and interference rates, I altered the strength of the relationship by increasing the magnitude of the slope parameters with a constant maximum or minimum and examined the resulting changes in the likelihood and strength of disruptive selection.

## Results

### Handling Times

When there is no correlation between attack rates and handling times, analysis of the model with a multispecies Holling Type-II functional response shows that increasing handling times reduce the parameter space over which disruptive selection occurs relative to consumers with a linear functional response (compare the dashed and solid lines in Figure 1A). Increasing handling times also reduce the strength of disruptive selection (Figure 1A). This decrease is due to relative changes in feeding rates across the consumer’s phenotypes (Figure 1B). In particular, as handling times increase, individuals with the highest attack rates show a decrease in feeding rates while individuals with lower attack rates show an increase in feeding rates (Figure 1B). This reduces the steepness of the fitness function at the fitness minimum thereby weakening disruptive selection. The relative changes in feeding rates among individuals with different phenotypes are the product of the interaction between attack rates, the saturating effect of handling times, and consequent changes in the equilibrium densities of resources (Figure 1B-D). As handling times increase, individuals with the highest attack rates saturate at increasingly lower resource densities (Figure 1C). The reduced feeding rates of individuals with the highest attack rates simultaneously increases equilibrium resource densities (Figure 1C-D). Individuals with low attack rates experience less saturation from handling times and thus show an increase in feeding rates due to the increase in equilibrium densities of the resources (Figure 1E).

**Figure 1:**
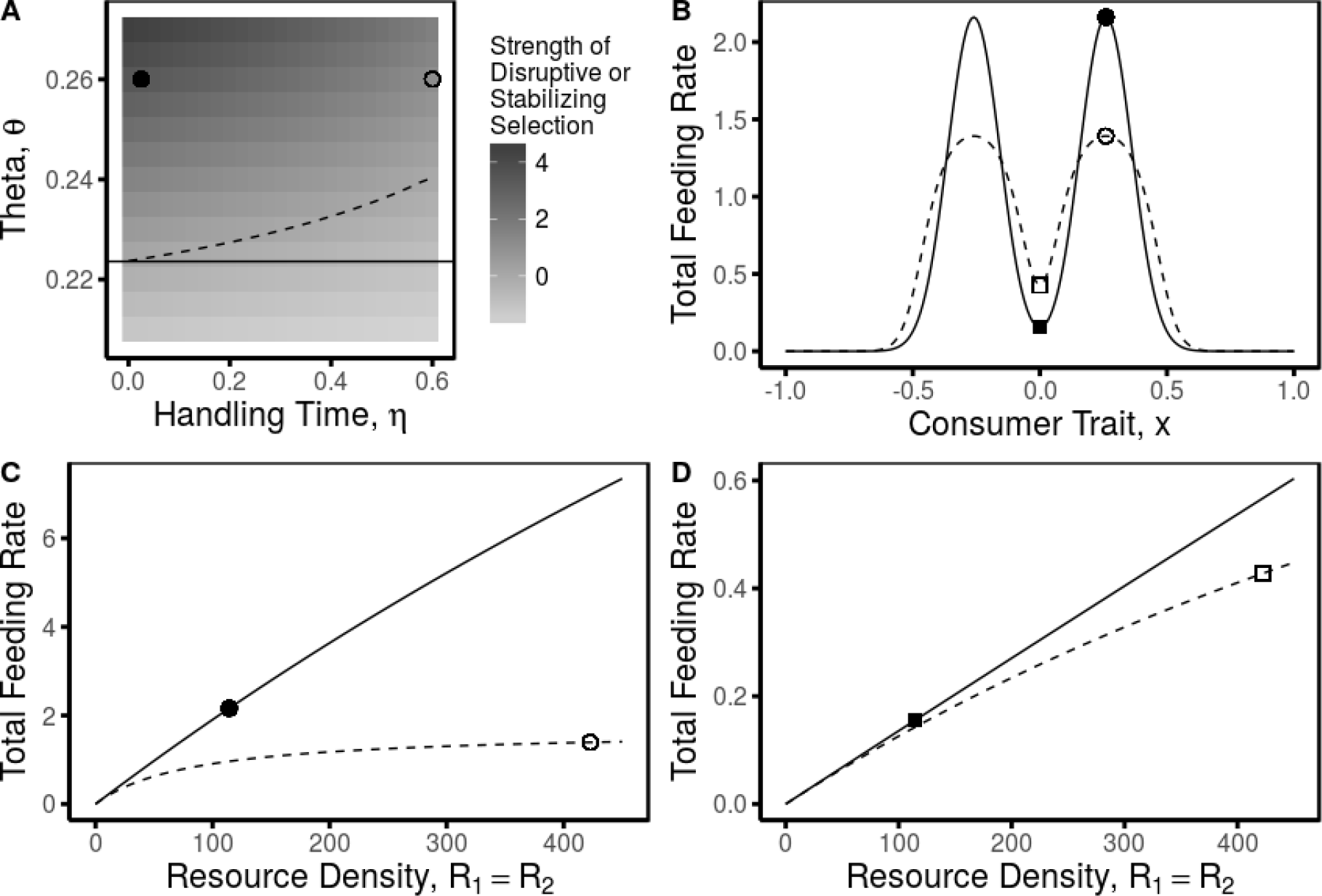
Larger handling times are associated with weaker disruptive selection (**A**). Larger handling times are also associated with a reduced parameter space over which disruptive selection occurs (above the dashed line in **A**) relative to the case in which consumers have linear functional responses (above the solid line in **A**). Handling times reduce the strength of disruptive selection by differentially affecting consumer feeding rates across the consumer’s phenotypes (compare feeding rates of consumers with low handling times in **B** (solid dot in **A**, solid lines in **B-D**) to feeding rates of consumers with high handling times (open dot in **A**, dashed lines in **B-D**)). For individuals with high attack rates (dots in **B** and **C**), feeding rates decrease with higher handling times despite an increase in equilibrium resource densities due to saturation caused by handling times (**C**). Individuals with low attack rates (squares in **B** and **D**) are less affected by the saturating effects of handling times and feeding rates increase with the increasing equilibrium resource densities (**D**). These changes in feeding rates reduce the potential gain in fitness associated with disruptive selection. Parameter used in the figure are: *x̄*, *σ* = 0.2, *α*_1,*max*_ = *α*_2,*max*_ = 0.02, τ_1_ = τ_2_ = 0.1, *r*_1_ = *r*_2_, *K*_1_ = *K*_2_ = 500, *e* = 0.5, *m* = 0.5.

The effects of handling time on the likelihood and strength of disruptive selection depend on whether ecological pleiotropy causes a positive or negative correlation between attack rates and handling times. If handling times and attack rates are positively correlated, then the parameter space over which disruptive selection occurs and the strength of disruptive selection decrease as the strength of the relationship increases (Figure 2A). In contrast, if handling times and attack rates are negatively correlated, the correlation weakens the effect of handling times on the likelihood and strength of disruptive selection (Figure 2C). Under certain parameter values, the negative correlation can increase the parameter space over which disruptive selection occurs relative to the case of linear consumer functional responses (Figure2C). These effects occur because correlations between attack rates and handling times either exacerbate or alleviate the saturating effects of handling times on individuals with high attack rates. When correlations between attack rates and handling times are positive, individuals with high attack rates experience greater saturation with an increase in the strength of the correlation causing a decreasing their feeding rates and increasing the feeding rates of consumers with low attack rates (Figure 2B). When correlations are negative, individuals with high attack rates experience less saturation with an increase in the strength of the correlation leading to higher feeding rates and decreases in the feeding rates of individuals with low attack rates (Figure 2D).

**Figure 2:**
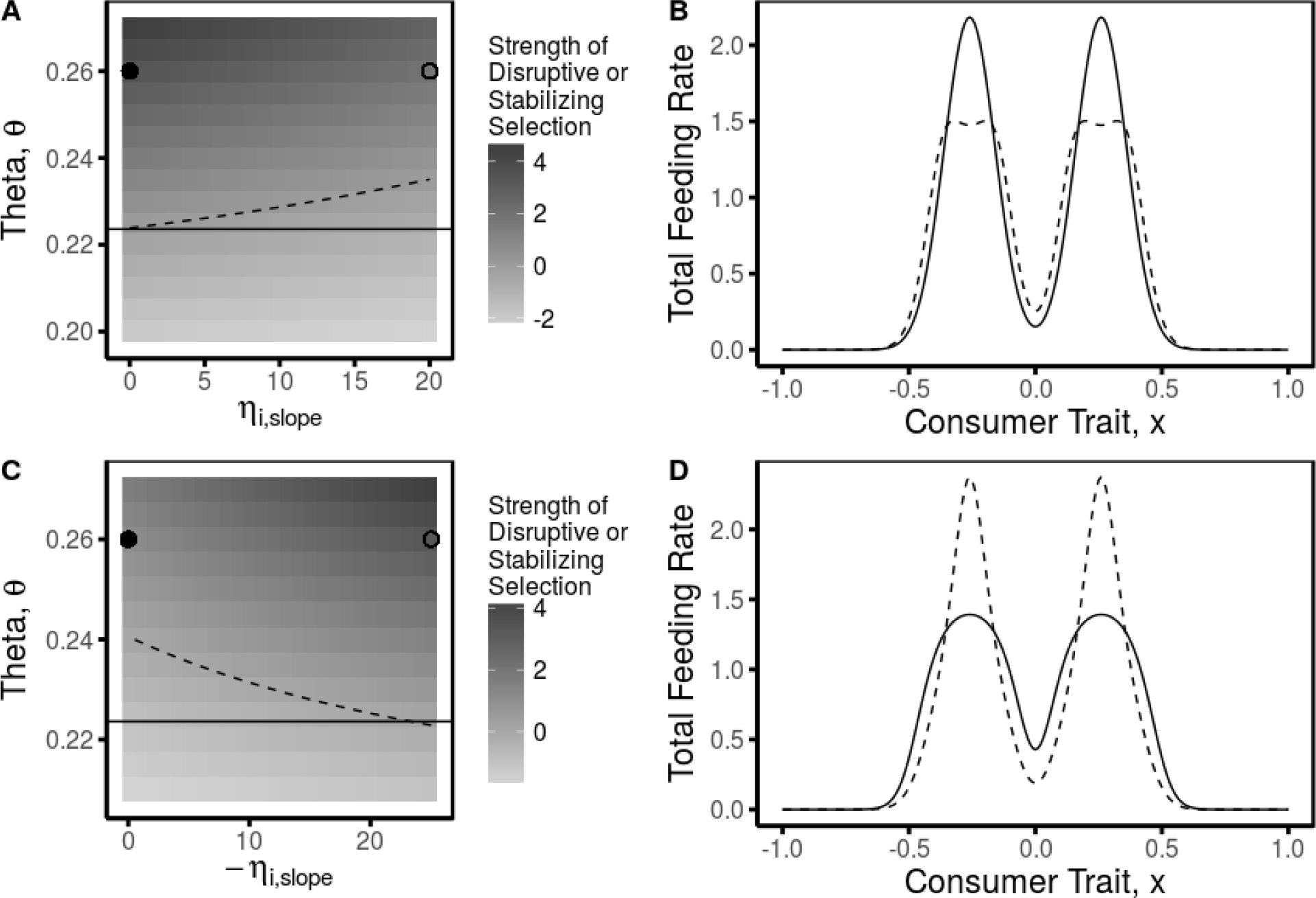
Ecological pleiotropy leading to correlations between attack rates and handling times alters the strength of disruptive selection (**A,C**). Positive correlations lead to a decrease in the strength of disruptive selection (**A**), while negative correlations lead to an increase in the strength of disruptive selection (**C**). The correlations also alter the parameter space over which disruptive selection occurs in this model (above the dashed lines in **A** and **C**) relative to the case in which consumers have linear functional responses (above the solid line in **A** and **C**). Changes in selection are due to changes across the consumer’s phenotypes in feeding rates as the correlation is changed from weak (solid dot in **A** and **C**, solid line in **B** and **D**) to strong (open dot in **A** and **C**, dashed line in **B** and **D**). Parameter values used in the figure are: *x̄*, *σ* = 0.2, *α*_1,*max*_ = *α*_2,*max*_ = 0.02, *θ*_1_ = −*θ*_2_ = 0.3, τ_1_ = τ_2_, *r*_1_ = *r*_2_ = 0.2, *K*_1_ = *K*_2_ = 500, *e* = 0.5, *m* = 0.5, and *η*_1,*min*_ = *η*_2,*min*_ = 0.01 in **A** and **B**, and *η*_1,*max*_ = *η*_2,*max*_ = 0.4 in **C** and **D**.

### Interference Rates

When there is no correlation between attack rates and interference rates, the effects of interference on the likelihood and strength of disruptive selection depend on the functional response considered. If interference is modeled using the Beddington-DeAngelis functional response, interference has no effect on the strength of disruptive selection (Figure 3A). As interference rates increase, the feeding rates across phenotypes remain constant because equilibrium resource and consumer densities change while feeding rates across phenotypes remain constant (Figure 3B). In contrast, if interference is modeled using the Crowley-Martin functional response, increasing interference decreases the parameter space over which disruptive selection occurs and decreases the strength of disruptive selection (Figure 3C). In contrast to the Beddington-DeAngelis functional response, interference interacts with the consumer’s attack rates and thus phenotype in the Crowley-Martin functional response and alters feeding rates across phenotypes (Figure 3D).

**Figure 3:**
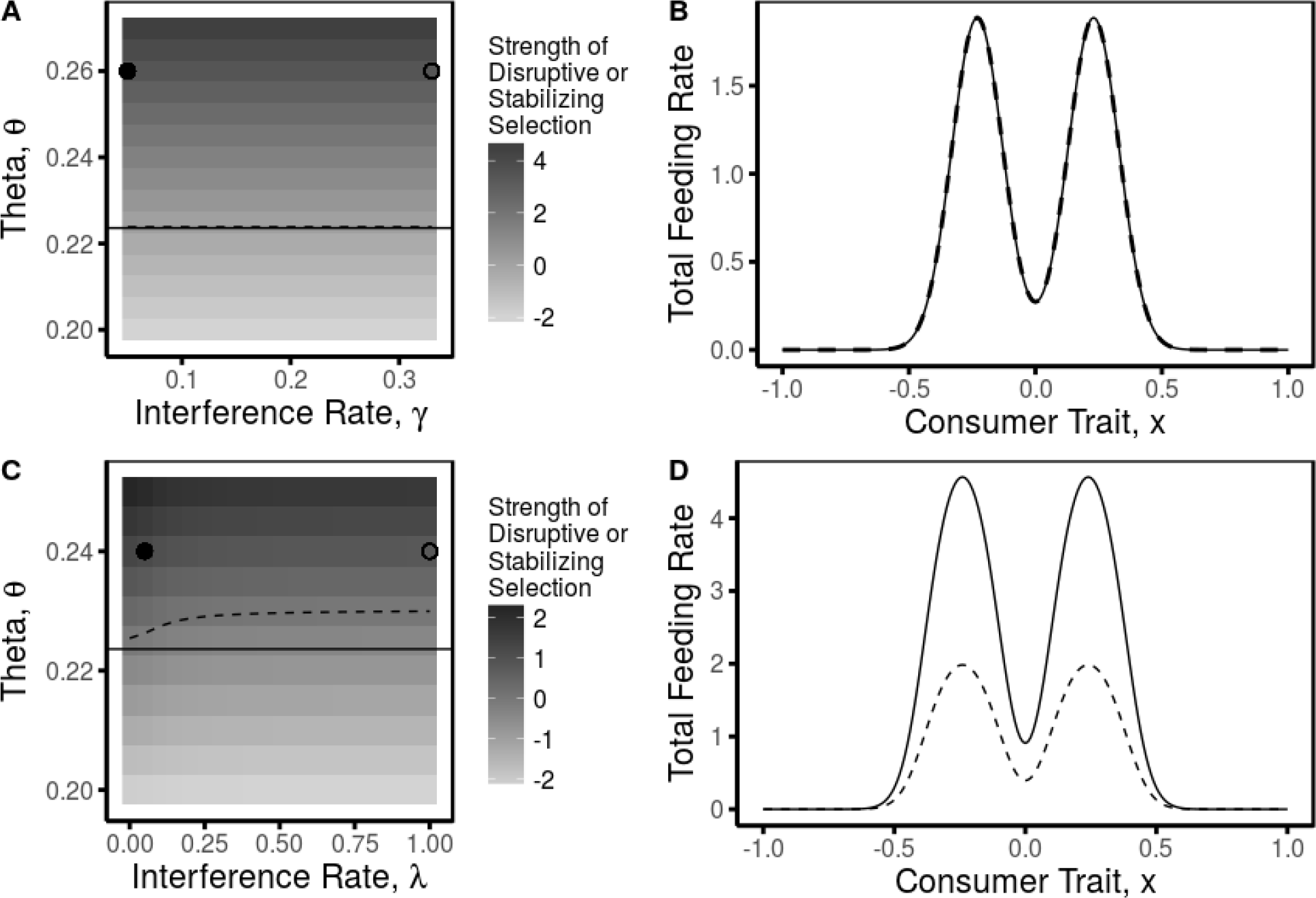
The effects of interference on the strength of disruptive selection are dependent on which functional response is considered (**A** – Beddington-DeAngelis functional response,**C** – Crowley-Martin functional response). The interference rate of the Beddington-DeAngelis functional response, (*γ*), has no effect on the strength of disruptive selection (**A**) and the parameter space over which disruptive selection occurs in this model (above the dashed line in **A**) compared to the case in which consumers have linear functional responses (above the solid line in **A**). This is because the interference rate but has no effect on the feeding rates of the consumers across their phenotypes (low interference: solid dot in **A**, solid line in **B**; high interference: open dot in **A**, dashed line in **B**). The interference rate of the Crowley-Martin functional response (λ) decreases the strength of disruptive selection (**C**) and parameter space over which disruptive selection occurs in this model (above the dashed line in **C**) relative to the model with linear consumer functional responses (above the solid line in **C**). The interference rate of the Crowley-Martin functional response differentially effects the feeding rates of consumers across the consumers phenotype (low interference: solid dot in **C**, solid line in **D**; high interference: open dot in **C** and dashed line in **D**). Parameter values used in the Beddington-DeAngelis figures are: *x̄* = 0, *σ* = 0.2, *α*_1,*max*_ = *α*_2,*max*_ = 0.3, *τ*_1_ = *τ* = 0.1 *r*_1_ = *r*_2_ = 0.2, *K*_1_ = *K*_2_ = 500, *e* = 0.5, *m* = 0.2, *η* = *η* = 0.01 Parameter values used in the Crowley-Martin figures are: *x̄* = 0, *σ* = 0.2, *α*_1,*max*_ = *α*_2,*max*_ = 0.1, *τ*_1_ = *τ* = 0.1 *r*_1_ = *r*_2_ = 0.2, *K*_1_ = *K*_2_ = 100, *e* = 0.5, *m* = 0.5, *η* = *η* = 0.1

As for handling times, the effects of interference on the likelihood and strength of disruptive selection are dependent on whether ecological pleiotropy causes correlations between attack rates and interference rates. Regardless of the functional response considered, a positive relationship between attack rate and interference leads to a decrease in the parameter space over which disruptive selection occurs and the strength of disruptive selection (Figures 4A,5A). A negative relationship between attack rates and interference leads to an increase in parameter space over which disruptive selection occurs and the strength of disruptive selection (4C,5C). These effects occur because the correlation causes the saturating effect of interference to affect consumers differently across phenotypes. For positive relationships between attack rates and interference, phenotypes with the highest attack rates experience the most saturation and reduced feeding rates, while this increases feeding rates for individuals with low attack rates because of increased equilibrium resource densities (Figures 4B,5B). For negative relationships between attack rates and interference, consumers with high attack rates experience less interference and have increased feeding rates, while consumers with low attack rates experience higher interference and lower equilibrium resource densities causing a decrease in feeding rates (Figures 4D,5D). For either functional response including interference, negative relationships between attack rates and interference are capable of increasing the parameter space in which selection is disruptive relative to linear functional responses (Figure 4C, 5C).

**Figure 4:**
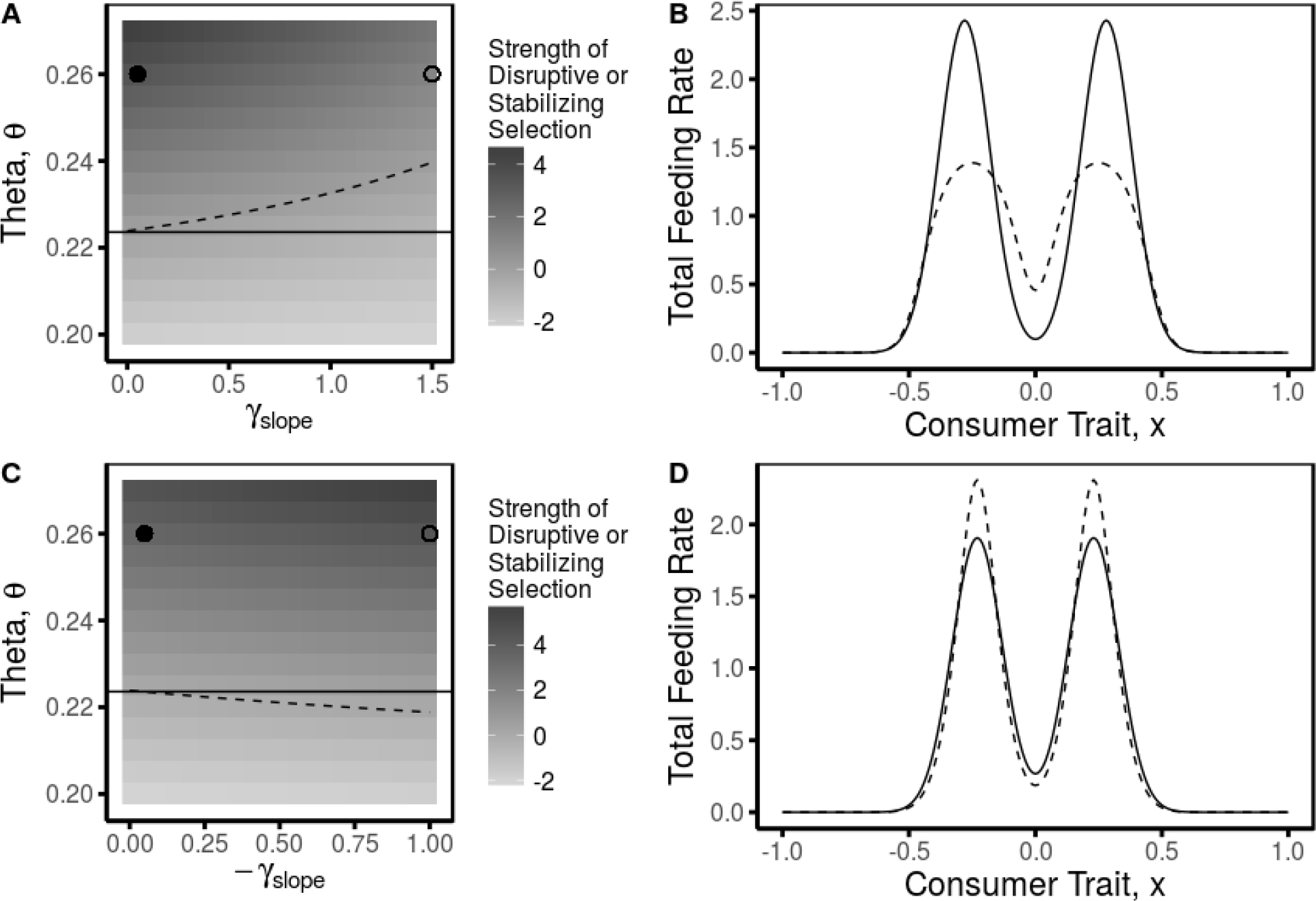
Ecological pleiotropy leading to correlations between attack rates and interference rates in the Beddington-DeAngelis functional response alters the strength of disruptive selection (**A**, **C**). Positive correlations lead to a decrease in the strength of disruptive selection (**A**), while negative correlations lead to an increase in the strength of disruptive selection (**C**). The correlations also alter the parameter space over which disruptive selection occurs in this model (above the dashed lines in **A** and **C**) relative to the case in which consumers have linear functional responses (above the solid line in **A** and **C**). Changes in selection are due to changes across the consumer’s phenotypes in feeding rates as the correlation is changed from weak (solid dot in **A** and **C**, solid line in **B** and **D**) to strong (open dot in **A** and **C**, dashed line in **B** and **D**). Parameter values used in the figure are: *x̄*, = 0 *σ* = 0.2, *α*_1,*max*_ = *α*_2,*max*_ = 0.3, *θ*_1_ = −*θ*_2_ = 0.3, τ_1_ = τ_2_, *r*_1_ = *r*_2_ = 0.2, *K*_1_ = *K*_2_ = 500, *e* = 0.5, *m* = 0.2, and *η*_1_ = *η*_2_ = 0.01, and in **A** and **B** *γ*_1,*min*_ = *γ*_2,*min*_ = 0.01 and in **C** and **D** *γ*1, *max γ*_2,*max*_ = 0.3.

**Figure 5:**
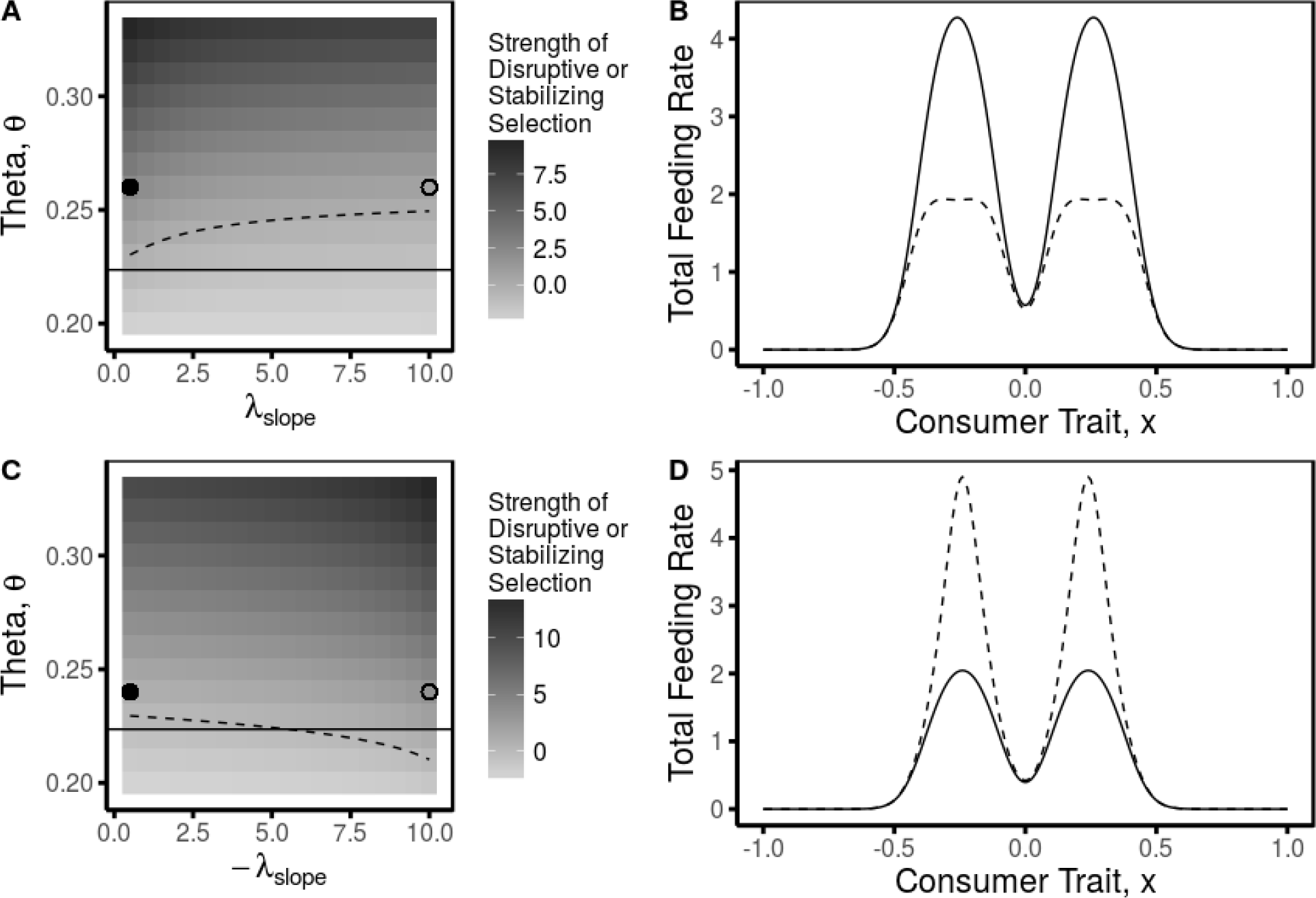
Ecological pleiotropy leading to correlations between attack rates and interference rates in the Crowley-Martin functional response alters the strength of disruptive selection (**A**, **C**). Positive correlations lead to a decrease in the strength of disruptive selection **A**, while negative correlations lead to an increase in the strength of disruptive selection (**C**). The correlations also alter the parameter space over which disruptive selection occurs in this model (above the dashed lines in **A** and **C**) relative to the case in which consumers have linear functional responses (above the solid line in **A** and **C**). Changes in selection are due to changes across the consumer’s phenotypes in feeding rates as the correlation is changed from weak (solid dot in **A** and **C**, solid line in **B** and **D**) to strong (open dot in **A** and **C**, dashed line in **B** and **D**). Parameter values used in the figure are: *x̄*, = 0 *σ* = 0.2, *α*_1,*max*_ = *α*_2,*max*_ = 0.1, τ_1_ = τ_2_ = 0.2, *r*_1_ = *r*_2_ = 0.2, *K*_1_ = *K*_2_ = 500, *e* = 0.5, *m* = 0.5, and *η*_1_ = *η*_2_ = 0.1, and in **A** and **B** *λ*_1,*min*_ = *λ*_2,*min*_ = 0.01 and in **C** and **D** *λ*1, *max* λ_2,*max*_ = 1.

## Discussion

The mechanisms generating and maintaining disruptive selection in generalist consumers have played a large role in our theoretical understanding of disruptive selection. However, predicting the circumstances under which disruptive selection in consumers is most likely and should be strongest remains difficult as theory on the ecological mechanisms altering ecological opportunity and intraspecific competition in consumers is still being developed. Here I show that two widespread factors influencing the strength of consumer resource interactions – nonlinear functional responses and ecological pleiotropy – are capable of altering the strength and likelihood of disruptive selection in consumers. These results support previous studies asserting that the strength of consumer-resource interactions can influence disruptive selection (Abrams and Ginzburg, 2000; Jones and Post, 2013, 2016) and suggest that disruptive selection is most likely in consumers whose traits not only determine their attack rates on resources but also reduce their handling times on those resources and their interference rates with other consumers.

The consumer-resource models presented here predict that disruptive selection in generalist consumers should be most common in populations in which handling times and interference are low or negatively correlated with attack rates. Confirmation of this with existing data is difficult, but some of the best known examples of disruptive selection in consumers do exhibit some of these features. For example, disruptive selection in beak size of the medium ground finch (*Geospiza fortis*) of the Galapagos Islands has been shown to be related to the ability of finches with different beak sizes to handle different sized seeds (Hendry et al., 2009; Schluter and Grant, 1984). Thus, attack rates and handling times should be negatively correlated with large-beaked individuals preferring and having lower handling times on large seeds and small-beaked individuals preferring and having lower handling times on small seeds. Another canonical example of disruptive selection is disruptive selection in several traits related to the use of benthic v. limnetic resources in three spine stickleback (Schluter, 1993). Morphological differences among individuals in resource-use traits have been shown to correlate with the feeding efficiency and growth rates of stickleback in benthic versus limnetic habitats (Schluter, 1993, 1995). How interference operates in this system is unclear, but the use of separate habitats may reduce interference by reducing the potential number of competitors or increase interference by concentrating individuals within habitats. Future empirical work should aim to more explicitly examine the relationship between consumer functional responses and natural selection. One possibility for doing so is to use comparative studies across populations or species measuring both functional responses and disruptive selection. Another possibility is to estimate the functional responses of individuals and correlate these functional responses to individual fitness proxies. Recent advances in estimating functional responses from observational data in the field (Novak et al., 2017) and the long history of estimating natural selection (Kingsolver et al., 2001; Lande and Arnold, 1983) should facilitate this effort.

The effects of handling time and interference on the strength of disruptive selection were largely dependent on whether the underlying trait was assumed to have pleiotropic effects that caused correlations between these parameters and the attack rates. The parameters describing nonlinear functional responses are likely to be correlated due to their determination by the same traits, however, the sign and strength of correlations among parameters are likely to be system specific. Some generalizations nevertheless might be possible. Optimal foraging theory, for example, suggests that individuals feeding on energetically equivalent resources should prefer the resources on which they have the lowest handling times (Stephens and Krebs, 1986). This will cause a negative correlation between attack rates and handling times among individuals (e.g. Tinker et al., 2007) which should increase the strength of disruptive selection. Positive correlations seem less likely. One possible source of this pattern could be changes in preferred resource size with consumer body size. For example, relative to mid-sized individuals, large bodied consumers may have higher attack rates on larger prey which require longer handling times, while small bodied consumers may have higher attack rates on smaller bodied resources but have higher handling times due to inefficiencies in handling resources given their size (Hassel et al., 1976).

Interference rates could have both positive or negative correlations with attack rates. For example, positive correlations have been observed in at least one protist system where both parameters were positively related to the swimming speed of the consumer (DeLong and Vasseur, 2013). However, the case could also be made that interference and attack rates should be negatively correlated. For example, attack rates, at least among species, commonly scale with body size (Berlow et al., 2009; Brose, 2010; Rall et al., 2012; Schneider et al., 2012), a trait that often confers an advantage in bouts of interference which could reduce the effects of interference on individuals with high attack rates (Rowland, 1989; Schoener, 1983). Individuals with high attack rates may also use resources more efficiently which may reduce their exposure to interference, at least in the Crowley-Martin functional response. Furthermore, individuals with high attack rates on particular resources may inhabit different habitats. Examples of this occur in several lake fish species, such as three-spined stickleback *(Gasterosteus aculeatus)* which have polymorphisms associated with using either benthic versus limnetic habitats (Lavin and McPhail, 1985; Schluter, 1995). As mentioned above, the use of different habitats may constrain the number of individuals that interfere with one another, thereby leading to an overall reduction in interference relative to the case in which all individuals use the same habitat. Conversely, individuals with high attack rates on the same resource may be concentrated within the same habitat, increasing interference. Unfortunately, data on the correlations among parameters in nonlinear functional responses are sparse, although traits that simultaneously influence multiple functional response parameters should be common.

To model the effects of mutual interference competition on consumer functional responses, I used two of the more ‘mechanistic’ functional response models including interference – the Beddington-DeAngelis and Crowley-Martin functional responses (Beddington, 1975; Crowley and Martin, 1989; DeAngelis et al., 1975). Which functional response was considered influenced the effects of interference on the strength of disruptive selection. The two functional responses differ primarily in whether or not interference occurs while consumers are handling resources. The most appropriate model for interference therefore is likely dependent on the biology of the particular system. For example, in systems with kleptoparasitism, fighting amongst consumers over already captured resources, or increased handling times in the presence of other consumers, the Crowley-Martin functional response may more appropriate (e.g. Ens and Goss-Custard, 1984; Norris and Johnstone, 1998; Smallegange et al., 2006; Zimmermann et al., 2015). In contrast, in systems where interference occurs largely separate from the handling of resources, or where time spent foraging is distinct from time spent interfering, the Beddington-DeAngelis functional response is likely to be more appropriate (e.g. Getty, 1981; Kratina et al., 2009; Pyke, 1979). Although the Crowley-Martin and Beddington-DeAngelis functional responses are sometimes unable to be distinguished statistically (Lang et al., 2012; Skalski and Gilliam, 2001; Stier and White, 2014; Zimmermann et al., 2015), the models here suggest that distinguishing among them mechanistically may be important to understand the evolutionary consequences of interference. Lastly, the functional responses used here assumed that interference rates, or their relationships with attack rates, were equal for both resources. Recent evidence has suggested that this may not be the case and that interference rates and facilitation effects may be prey-specific in nature (Novak et al., 2017). These results suggest that theory incorporating prey-specific interference rates may be needed to understand the ecological and evolutionary consequences of interference with more than one resource.

There are several potential outcomes of disruptive selection and the particular outcome may be a function of the strength of disruptive selection (Rueffler et al., 2006). Under the quantitative genetics framework, the most likely outcome of disruptive selection is an increase in the phenotypic variation of the consumer’s resource-use trait. Patel and Schreiber (2015) have shown that the second derivative of the fitness function used here to measure disruptive and stabilizing selection also determines selection at the evolutionary equilibrium under the adaptive dynamics approach for modeling evolution. Under the adaptive dynamics framework, this theory would predict that evolutionary branching and speciation would occur at the fitness minimum. In general, the outcome of disruptive selection is likely to depend on system specific factors such as the underlying genetics and mating system of the population in concert with the strength of selection. For example, although ecological opportunity and intraspecific resource competition may cause disruptive selection on a trait, opposing directional or stabilizing selection on that trait from other sources, or gene flow, might overwhelm weak disruptive selection (Lande and Arnold, 1983; Nosil, 2012). Future theory explicitly examining how the strength of selection is likely to lead to different evolutionary outcomes will help to further understand how changes to the strength of disruptive selection through ecological factors are likely to manifest in nature.

Overall, my results support the assertions of Abrams et al. (2008) and Jones and Post (2013, 2016) that the strength and likelihood of disruptive selection is dependent on the strength of the underlying consumer-resource interactions. In the model presented here, the effects of nonlinear functional responses and pleiotropy are largely due to how handling resources or interference produces saturation in the feeding rates of the consumers with the highest attack rates. If saturation is increased, the consumer-resource interactions weaken resulting in weakened disruptive selection and vice versa. Abrams et al. (2008) and Jones and Post (2013, 2016), however, have predicted a unimodal relationship between the strength of consumer-resource interactions and the strength of disruptive selection. Their reasoning is that when consumer-resource interactions are very weak there is little depletion of resources, intraspecific resource competition is weak, and thus disruptive selection is weak. In contrast, when consumer-resource interactions are very strong resources associated with the most common consumer phenotypes will go locally extinct, ecological opportunity will decrease, and disruptive selection will weaken. Allowing for this effect would require extending the theory here to models with a continuous distribution of resources such as the MacArthur model which has been used widely to model disruptive selection in consumers (MacArthur, 1972). Caution should be taken in doing so. The MacArthur model with linear functional responses has been shown to have a stable global equilibrium (Chesson, 1990). Nonlinear functional responses, however, can lead to cycling in population dynamics which may alter evolutionary dynamics (Svanbäck et al., 2009). Nevertheless, the incorporation of nonlinear functional responses and pleiotropy into these models would be worthwhile.

Lastly, consumer-resource models similar to those used to examine disruptive selection have also been used to study other topics such as the coevolution of competitors and character displacement (Case, 1981; Doebeli, 1978; Roughgarden, 1976; Taper and Case, 1992). As I have shown here that nonlinear functional responses and ecological pleiotropy can alter the likelihood and strength of disruptive selection, these factors may also influence phenomena such as character displacement. Previous studies have also suggested as much. For example, Abrams (1980), has shown that nonlinear functional responses alter the strength of competition among consumers in a two consumer-two resource system. Further incorporation of nonlinear functional responses and ecological pleiotropy into evolutionary theory will provide insight into how these widespread ecological factors influence evolutionary dynamics beyond disruptive selection.

## Supporting information

## Literature Cited

Abrams, P. A. 1980. Consumer functional response and competition in consumer-resource systems. Theoretical Population Biology 17:80–102.

Abrams, P. A., and L. R. Ginzburg. 2000. The nature of predation: prey dependent, ratio dependent, or neither? Trends in Ecology and Evolution 15:337–341.

Abrams, P. A., Y. Harada, and H. Matsuda. 1993. On the relationship between quantitative genetic and ess models. Evolution 47:982–985.

Abrams, P. A., C. Rueffler, and G. Kim. 2008. Determinants of the strength of disruptive and/or divergent selection arising from resource competition. Evolution 62:1571–1586.

Ackermann, M., and M. Doebeli. 2004. Evolution of niche width and adaptive diversification. Evolution 58:2599–2612.

Barbour, M. A., M. A. Fortuna, J. Bascompte, J. R. Nicholson, R. Julkunen-Tiitto, E. S. Jules, and G. M. Crutsinger. 2016. Genetic specificity of a plant-insect food web: Implications for linking genetic variation to network complexity. Proceedings of the National Academy of Sciences page 201513633.

Beddington, J. R. 1975. Mutual interference between parasites or predators and its effect on searching efficiency. Journal of Animal Ecology 51:331–340.

Berlow, E. L., J. A. Dunne, N. D. Martinez, P. B. Stark, R. J. Williams, and U. Brose. 2009. Simple prediction of interaction strengths in complex food webs. Proceedings of the National Academy of Sciences 106:187–191.

Bolnick, D. I. 2001. Intraspecific competition favours niche width expansion in Drosophila melanogaster. Nature 410:463–466.

Bolnick, D. I. 2004. Can intraspecific competition drive disruptive selection? An experimental test in natural populations of sticklebacks. Evolution 58:608–618.

Bolnick, D. I., P. Amarasakare, M. S. Araujo, R. Burger, J. M. Levine, M. Novak, V. H. Rudolf, S. J. Schreiber, M. C. Urban, and D. Vasseur. 2011. Why intraspecific trait variation matters in community ecology. Trends in Ecology and Evolution 26:183–192.

Bolnick, D. I., and O. L. Lau. 2008. Predictable patterns of disruptive selection in stickleback in postglacial lakes. The American Naturalist 172:1–11.

Brose, U. 2010. Body-mass constraints on foraging behavior determine population and food-web dynamics. Functional Ecology 24:28–34.

Case, T. J. 1981. Niche packing and coevolution in competition communities. Proceedings of the National Academy of Sciences USA 78:5021–5025.

Chesson, P. 1990. Macarthur’s consumer-resource model. Theoretical Population Biology 37:26–38.

Crowley, P. H., and E. K. Martin. 1989. Functional responses and interference within and between year classes of a dragonfly population. Journal of the North American Benthological Society 8:211–221.

DeAngelis, D. L., R. A. Goldstein, and R. V. O’Neill. 1975. A model for trophic interaction. Ecology 56:881–892.

DeLong, J. P. 2017. Ecological pleiotropy suppresses the dynamic feedback generated by a rapidly changing trait. The American Naturalist 189:592–597.

DeLong, J. P., and J. P. Gibert. 2016. Gillespie eco-evolutionary models (GEMs) reveal the role of heritable trait variation in eco-evolutionary dynamics. Ecology and Evolution 6:935–945.

DeLong, J. P., and D. A. Vasseur. 2011. Mutual interference is common and mostly intermediate in magnitude. BMC Ecology 11:1.

DeLong, J. P., and D. A. Vasseur 2013. Linked exploitation and interference competition drives the variable behavior of a classic predator-prey system. Oikos 122:1393–1400.

Dieckmann, U., and M. Doebeli. 1999. On the origin of species by sympatric speciation. Nature 400:354–357.

Doebeli, M. 1978. Competitive speciation. Biological Journal of the Linnean Society 10:275–289.

Doebeli, M. 2011. Adaptive Diversification. Princeton University Press, New Jersey.

Ens, B. J., and J. D. Goss-Custard. 1984. Interference among oystercatchers Haematopus ostralegus, feeding on mussels, Mytilus edulis, on the exe estuary. Journal of Animal Ecology 53:217–231.

Getty, T. 1981. Territorial behavior of eastern chipmunks (Tamias striatus): Encounter avoidance and spatial time-sharing. Ecology 62:915–921.

Gibert, J. P., and C. E. Brassil. 2014. Individual phenotypic variation reduces interaction strengths in a consumer-resource system. Ecology and Evolution 4:3703–3713.

Gibert, J. P., and J. P. DeLong. 2017. Phenotypic variation explains food web structural patterns. Proceedings of the National Academy of Sciences page 201703864.

Hart, S. P., S. J. Schreiber, and J. M. Levine. 2016. How variation between individuals affect species coexistence. Ecology Letters 19:825–838.

Hassel, M. P., J. H. Lawton, and J. R. Beddington. 1976. The components of arthropod predation: I. The prey death-rate. Journal of Animal Ecology 45:135–164.

Hassel, M. P., and G. C. Varley. 1969. New inductive population model for insect parasites and its bearing on biological control. Nature 223:1133–1137.

Hendry, A. P., S. K. Huber, L. F. De León, A. Herrel, and J. Podos. 2009. Disruptive selection in a bimodal population of darwin’s finches. Proceedings of the Royal Society B 276:753–759.

Holling, C. S. 1959. The components of predation as revealed by a study of small mammal predation of the european pine sawfly. The Canadian Entomologist 91:293–320.

Hughes, R. A., B. D. Inouye, M. T. Johnson, N. Underwood, and M. Vellend. 2008. Ecological consequences of genetic diversity. Ecology Letters 11:609–623.

Iwasa, Y., A. Pomiankowski, and S. Nee. 1991. The evolution of costly mate preferences ii. the “handicap” principle. Evolution 45:1431–1442.

Jeschke, J. M., M. Kopp, and R. Tollrian. 2004. Consumer-food systems: why type I functional responses are exclusive to filter feeders. Biological Reviews 79:337–349.

Jones, A. J., and D. M. Post. 2013. Consumer interaction strength may limit the diversifying effect of intraspecific competition: a test in alewife (Alosa pseudoharengus). The American Naturalist 181:815–826.

Jones, A. J., and D. M. Post. 2016. Does intraspecific competition promote variation? a test via synthesis. Ecology and Evolution 6:1646–1655.

Kingsolver, J. G., H. E. Hoekstra, J. M. Hoekstra, D. Berrigan, S. N. Vigniere, C. E. Hill, A. Hoang, P. Gibert, and P. Beerli. 2001. The strength of phenotypic selection in natural populations. The American Naturalist 157:245–261.

Kratina, P., M. Vos, A. Bateman, and B. R. Anholt. 2009. Functional responses modified by predator density. Oecologia 159:425–433.

Lande, R. 1976. Natural selection and random genetic drift in phenotypic evolution. Evolution 30:314–334.

Lande, R., and S. J. Arnold. 1983. The measurement of selection on correlated characters. Evolution 37:1210–1226.

Lang, B., B. C. Rall, and U. Brose. 2012. Warming effects of consumption and intraspecific interference competition depend on predator metabolism. Journal of Animal Ecology 81:516523.

Lavin, P. A., and J. D. McPhail. 1985. The evolution of freshwater diversity in the threespine stickleback (Gasterosteus aculeatus): Site-specific differentiation of trophic morphology. Canadian Journal of Zoology 63:2632–2638.

Lawlor, L. R., and J. M. Smith. 1976. The coevolution and stability of competing species. The American Naturalist 110:79–99.

MacArthur, R. H. 1972. Geographical Ecology. Princeton University Press, New Jersey.

Martin, R. A., and D. W. Pfennig. 2009. Disruptive selection in natural populations: The roles of ecological specialization and resource competition. The American Naturalist 147:268–281.

Martin, R. A., and D. W. Pfennig. 2012. Widespread disruptive selection in the wild is associated with intense resource competition. BMC evolutionary biology 12:136.

Norris, K., and I. Johnstone. 1998. Interference competition and the functional response of oys-tercatchers searching for cockles by touch. Animal Behaviour 56:639–650.

Nosil, P. 2012. Ecological Speciation. Oxford University Press, New York.

Novak, M., C. Wolf, K. E. Coblentz, and I. D. Shepard. 2017. Quantifying predator dependence in the functional response of generalist predators. Ecology Letters 20:761–769.

Patel, S., and S. J. Schreiber. 2015. Evolutionarily driven shifts in communities with intraguild predation. The American Naturalist 186:E98–E110.

Pyke, G. H. 1979. The economics of territory size and time budget in the golden-winged sunbird. The American Naturalist 114:131–145.

Rall, B. C., U. Brose, M. Hartvig, G. Kalinkat, F. Schwarzmüller, O. Vucic-Pestic, and O. L. Petchey. 2012. Universal temperature and body-mass scaling of feeding rates. Phil. Trans. R. Soc. B 367:2923–2934.

Roughgarden, J. 1976. Resource partitioning among competing species: coevolutionary approach. Theoretical Population Biology 9:388–424.

Rowland, W. J. 1989. The effects of body size, aggression and nuptial coloration on competition for territories in male threespine sticklebacks, Gasterosteus aculeatus. Animal Behaviour 37:282–289.

Rueffler, C., T. J. M. Van Dooren, O. Leimar, and P. A. Abrams. 2006. Disruptive selection and then what? Trends in Ecology and Evolution 238-245:238–345.

Schluter, D. 1993. Adaptive radiation in sticklebacks: Size, shape, and habitat use efficiency. Ecology 74:699–709.

Schluter, D. 1995. Adaptive radiation in sticklebacks: trade-offs in feeding performance and growth. Ecology 76:82–90.

Schluter, D. 2000. The ecology of adaptive radiation. Oxford University Press, Oxford.

Schluter, D., and P. R. Grant. 1984. Ecological correlates of morphological evolution in a Darwin’s finch Geospiza difficilis. Evolution 38:856–869.

Schneider, F. D., S. Scheu, and U. Brose. 2012. Body mass constraints on feeding rates determine the consequences of predator loss. Ecology Letters 15:436–443.

Schoener, T. W. 1983. Field experiments on interspecific competition. The American Naturalist 122:240–285.

Schreiber, S. J., R. Burger, and D. I. Bolnick. 2011. The community effects of phenotypic and genetic variation within a predator population. Ecology 92:1582–1593.

Schreiber, S. J., S. Patel, and C. terHorst. 2018. Evolution as a coexistence mechanism: Does genetic architecture matter? The American Naturalist 191:407–420.

Skalski, G. T., and J. F. Gilliam. 2001. Functional responses with predator interference: Viable alternatives to the holling type II model. Ecology 82:3083–3092.

Smallegange, I. M., J. van der Meer, R. H. J. M. Kurvers, and L. Persson. 2006. Disentangling interference competition for exploitative competition in a crab-bivalve system using a novel experimental approach. Oikos 113:157–167.

Smith, J. M. 1962. Disruptive selection, polymorphism and sympatric speciation. Nature 195:60–62.

Stephens, D. W., and J. R. Krebs. 1986. Foraging Theory. Princeton University Press, New Jersey.

Stier, A. C., and J. W. White. 2014. Predator density and the functional responses of coral reef fish. Coral Reefs 33:235–240.

Strauss, S. Y., and R. E. Irwin. 2004. Ecological and evolutionary consequences of multispecies plant-animal interactions. Annual Review of Ecology, Evolution, and Systematics 35:435–466.

Svanbäck, R., M. Pineda-Krch, and M. Doebeli. 2009. Fluctuating population dynamics promotes the evolution of phenotypic plasticity. The American Naturalist 174:176–189.

Svanbäck, R., M. Quevedo, J. Olsson, and P. Eklöv. 2015. Individuals in food webs: the relationships between trophic position, omnivory and among-individual diet variation. Oecologia 178:103–114.

Taper, M. L., and T. J. Case. 1992. Models of character displacement and the theoretical robustness of taxon cycles. Evolution 46:317–333.

Tinker, M. T., G. Bentall, and J. A. Estes. 2007. Food limitation leads to behavioral diversification and dietary specialization in sea otters. Proceedings of the National Academy of Sciences 105:560–565.

Vucic-Pestic, O., B. C. Rall, G. Kalinkat, and U. Brose. 2009. Allometric functional response model: body masses constrain interaction strengths. Journal of Animal Ecology 79:249–256.

Zimmermann, B., H. Sand, P. Wabakken, O. Liberg, and H. P. Anreassen. 2015. Predator-dependent functional response in wolves: from food limitation to surplus killing. Journal of Animal Ecology 84:102–112.622

